# Internal water storage buffering maintains plant function under drought as described by a general hydraulic model

**DOI:** 10.1101/2020.02.11.943563

**Authors:** Avigail Kaner, Yakir Preisler, José M. Grünzweig, Yair Mau

## Abstract

- Internal water storage is of crucial importance for plants under drought stress, allowing them to temporarily maintain transpiration higher than root-uptake flow, thus potentially keeping a positive carbon balance. A deep understanding of this adaptation is key for predicting the fate of ecosystems subjected to climate change-induced droughts of increasing intensity and duration.
- Using a minimalistic model, we derive predictions for how environmental drivers (atmospheric demand and soil water availability) interplay with the water storage, creating time lags between the flows in the plant, and granting the plant increased hydraulic safety margin protecting its xylem from embolism.
- We parametrize our model against transpiration and sap flow measurements in a semi-arid pine forest during seasonal drought. From the parametrized whole-stand traits, we derive a 3.7-hour time lag between transpiration and sap flow, and that 31% of daily transpiration comes directly from the plant’s internal water storage, both corroborated by the measurements.
- Due to the model simplicity, our results are useful for interpreting, analyzing, and predicting the effects of the internal storage buffering from the individual plant to the ecosystem scale. Because internal storage produces survival-enhancing behavior in sub-daily time scales, it is an indispensable component for modeling ecosystems under drought stress.

## Introduction

A plant’s internal water storage can act as a water buffer, decoupling transpiration flow from commensurate root-uptake flow. This buffering gives the plant crucial leeway in supporting transpiration and photosynthesis in times of greater drought stress, either from the soil or from the atmosphere.

The footprint of the water buffer offered by the internal water storage is felt throughout the whole plant-water dynamics. As a result of the water buffer (i) sap flow lags behind transpiration flow (Goldstein et al., 1998; Schäfer et al., 2000; Phillips et al., 2003; Kumagai et al., 2009); (ii) plant tissues responsible for holding the internal water storage expand and contract on a daily basis (Sevanto et al., 2002; Steppe et al., 2015); (iii) xylem water potential is granted a safety margin from very low values, decreasing embolism risk (Meinzer et al., 2009; Scholz et al., 2011; Oliva Carrasco et al., 2014); (iv) upon sudden changes in soil or atmospheric conditions, plant flows (transpiration, sap flow) respond with a characteristic relaxation time that is dependent on the internal water storage properties (Daley et al., 2008).

While there is a wealth of experimental evidence accounts for the internal water storage’s impact on plant hydraulics (Tyree and Yang, 1990; Holbrook and Sinclair, 1992; Holbrook, 1995; Meinzer et al., 2003; Scholz et al., 2011; Köcher et al., 2013), a deep understanding of the causal underpinnings between the internal storage and the effects mentioned above is still lacking. Many numerical models use internal water storage units as part of their formulation, with varying degrees of complexity and required parametrization (Cowan, 1972; Sperry et al., 1998; Ogee et al., 2003; Steppe et al., 2006; Bonan et al., 2014; Mirfenderesgi et al., 2016; Hartzell et al., 2017). These models succeed in capturing the effects produced by the internal water storage in the plant hydraulics, but due to the large number of mechanisms and parameters included in them, it can be cumbersome or impractical to tease out *causal relationships* and general trends in behavior produced by those mechanisms.

It is vital to expand our understanding of the role of the internal water storage in plant survival, as ecosystems around the world increasingly experience drought stress. Climate change is expected to intensify regional drying in the sub-tropics and in the Amazon, due to a combined increase in evaporative demand and decrease in precipitation (Neelin et al., 2006; Cook et al., 2014). The resilience of drought-stressed ecosystems might be contingent on their ability to leverage the sub-daily water dynamics produced by internal water storage buffering effects.

Our goal in this paper is to thoroughly examine the buffering mechanism offered by the internal water storage, and to quantify its impact on the dynamics of water flow throughout the plant. The results we derive regarding these dynamics are instrumental in determining whole-plant traits from measured water flows. We approach this goal by formulating the simplest possible model of plant hydraulics, which includes the internal water storage, and is driven by the environment through the soil and atmosphere. This model is amenable to the methods of system dynamics, which provides a powerful machinery to investigate the response of a system to arbitrary external forcing. Focusing on the plant response to periodic and step-like changes of the soil and leaf water potentials, we derive typical time scales of reaction, the time lag between daily peaks in transpiration and sap flow, and frequency filtering offered by the buffer effect.

It is important to make clear that we do not seek to build a comprehensive model for plant hydraulics. We focus here on one process only, namely the role of internal water storage, and ask: how much of the plant hydrodynamics can be attributed to it? As we show in the model evaluation section, this approach is robust even under the model assumption of constant stomatal conductance.

## Materials and Methods

### The hydraulic system and its electric analogue

Our starting point is the definition of a minimal hydraulic model for water flow in a plant, whose internal water storage plays an important role in the dynamics of water flows. The diagram in Fig. 1a represents our minimal model [similar to that of (Wronski et al., 1985; Katerji et al., 1986; Carlson and Lynn, 1991)], where water flows upwards, from the soil (bottom) to the leaves (top). Water potential is denoted by *ψ* (MPa), water flow rate is denoted by *Q* (mmol h^*−*1^), and water flow resistance is denoted by *R* (MPa h mmol^*−*1^).

**Figure 1:**
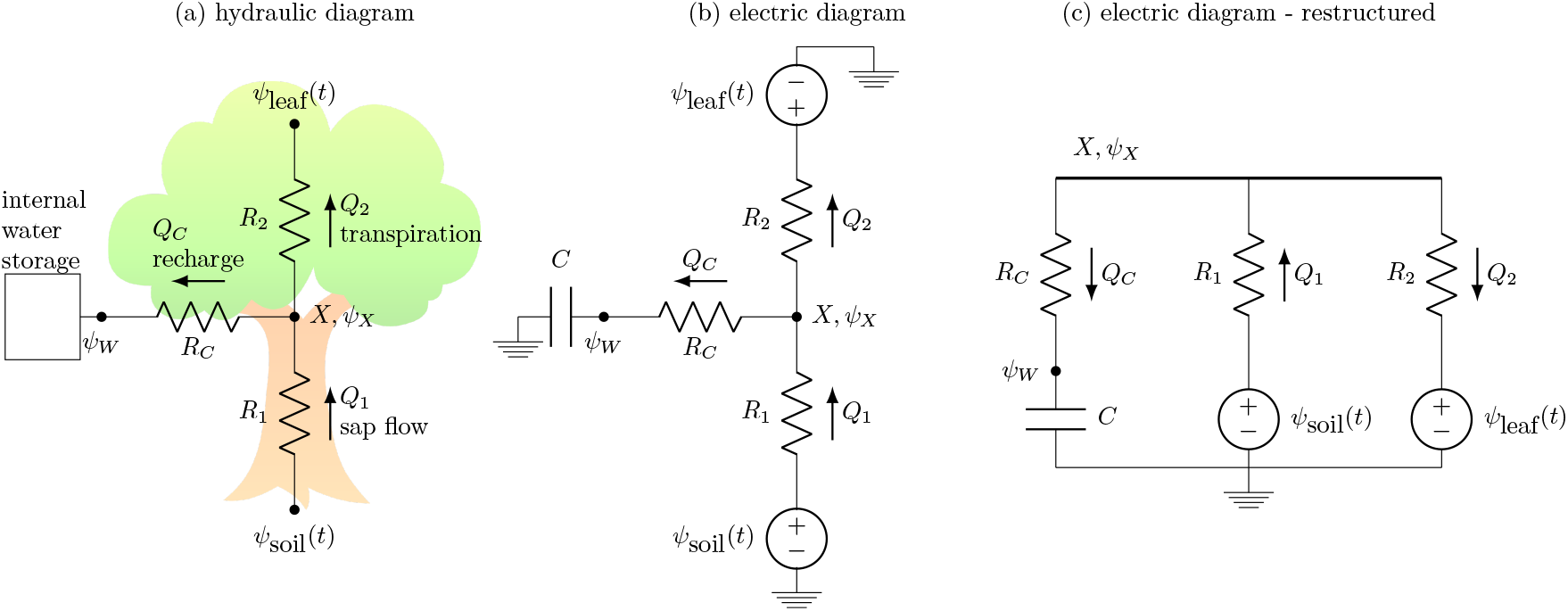
Diagram of the electric analogue gives deeper insight into how model components interact. Diagram a: minimal hydraulic model. Diagrams b, c: electric analogue of hydraulic model. All diagrams give the same dynamics, i.e., they are all equivalent.

We assume that a water storage unit of capacitance *C* (mmol MPa^*−*1^), representing the continuous water distribution throughout the plant’s various tissues, is connected to the xylem at node *X*. The water potential of the storage unit is called *ψ*_*W*_, and the water potential at *X* is called *ψ*_*X*_. The plant water dynamics are driven by two time-dependent external potentials, the soil water potential *ψ*_soil_(*t*) and the leaf water potential *ψ*_leaf_(*t*). As a result of these external potentials, water flows between the different nodes of the diagram, identified here by: sap flow *Q*_1_, transpiration flow *Q*_2_, and water storage recharge flow *Q*_*c*_. These flows have a Darcy-like (linear) dependence on water potential difference, and we call the resistances to flow *R*_1_, *R*_2_ and *R*_*c*_, respectively. In this model the stomatal conductance is fixed, therefore the leaf potential simply tracks atmospheric potential. The basic simplifying assumption of this model is that the resistances *R*_1_, *R*_2_, *R*_*c*_ and the hydraulic capacitance *C* of the water storage unit are constant (these quantities are discussed in greater detail below). Water fluxes *q* = *Q/A* (mmol h^*−*1^ m^*−*2^) could be used instead of water flows *Q*, where *A* is a unit area of soil. In that case, however, resistances and capacitances would need to be exchanged for resistivities and capacitivities (Hunt et al., 1991).

It can be useful to translate this basic hydraulic model into its electric analogue. This allows us to look at our problem from another point of view, and as we will see, it brings about new insights on the structure and behavior of the original system and on the fundamental assumptions regarding the hydraulic system.

We call two models analogue if the same set of equations can be used to describe them, and therefore they have the same dynamics. In a mathematical parlance, the two systems are called isomorphic, i.e., there is a set of translation rules from the hydraulic to the electric system that preserves the dynamics. We discuss below four rules needed to translate the diagram in Fig. 1a to the electric analogue description of Fig. 1b.

#### Rule 1: flow drivers

We assume a Darcy-like saturated flow, where the flow *Q* between two points in the plant is proportional to the water potential difference ∆*ψ* between them, according to

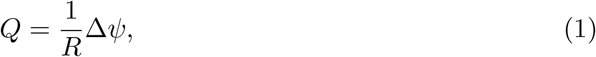

where the proportionality factor is the hydraulic conductance *K* = 1/*R*, and R is the resistance to water flow. This equation is analogous to Ohm’s law (van den Honert, 1948; Richter, 1973), ∆*V* = *RI*, where the current *I* is driven by a difference in electric potential ∆*V*, and *R* denotes the resistance to electric current. Both hydraulic and electric formulations could account for resistances *R*(*ψ*) that depend on the potential. In plant hydraulics, an increase in resistance (or loss in conductance) arises from xylem embolism as the water potential decreases. In this paper, however, we will consider *R* to be constant, assuming that xylem water potential does not reach low enough values conducive to embolism.

#### Rule 2: flow conservation

In both hydraulic and electric systems, water and electric charge conservation implies the conservation of flow. Kirchhoff’s Current Law of electricity is therefore analogous to

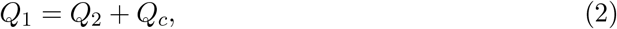

where the water flows *Q* meet at the node X in Fig. 1a. There is freedom to determine if positive *Q*_*c*_ means recharging or depleting the water storage, and in this paper *Q*_*c*_ denotes recharge, i.e., the water storage is filled for *Q*_*c*_ > 0.

#### Rule 3: external potentials and their reference points

All values of water potential *ψ* in hydraulics are implicitly reported with respect to an agreed-upon zero reference potential, which is set at water surface. In electric circuits, it is common to *explicitly* mark the zero electric potential using the ‘ground’ symbol. (In this paper *ground* refers exclusively to the zero electric potential, while ‘soil’ refers to the actual soil water potential.)

Because we treat the soil water potential *ψ*_soil_ and the leaf water potential *ψ*_leaf_ as being external drivers, in the electric analogue they are represented as time-dependent potential sources, with an explicit connection to the ground potential (see top and bottom extremes of Fig. 1b).

#### Rule 4: storage/capacitor

The water potential *ψ*_*W*_ of the storage unit is dependent on the water content *W* (mmol) in it, according to the desorption curve *ψ*_*W*_ = *f* (*W*) (also called pressure-volume curve). The capacitance *C* (mmol MPa^*−*1^) of the storage unit describes how strongly the plant tissues hold the water in them, given by

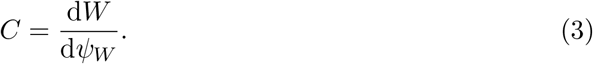

The electrical analogue of a water storage unit is the capacitor. Unlike a water storage unit, however, a capacitor is a two-terminal component, i.e., it has two wires that must be connected to the circuit. One connection is to the point marked as *ψ*_*W*_, while the other is grounded, as shown in Fig. 1b. This wiring of the capacitor means that the potential *ψ*_*W*_ is the potential across the capacitor, and it is explicitly reported in relation to the zero potential, in the same way done with the external drivers (see rule 2 above). This configuration is an exact analogue of the water storage unit shown in Fig. 1a, with the difference that there the water potential *ψ*_*W*_ is *implicitly* reported with respect to the zero potential.

In this paper we will consider *C* to be constant, although a varying capacitance could be used in either hydraulic or electric formulation according to a nonlinear desorption curve *f* (*ψ*). The underlying assumption behind a constant *C* value is that the time scales of the dynamics studied here (a few hours) are much shorter than the time it takes for the capacitance to significantly change due to water depletion of the plant tissues that form the internal storage (Hunt et al., 1991).

### The minimal model

The rules defined above allow us to convert the hydraulic system of Fig. 1a into its electric analogue shown in Fig. 1b. If both hydraulic and electric descriptions are equivalent, what do we gain by translating the model from one formulation into the other? Figure 1c shows the exact same electric diagram as in Fig. 1b, but with some restructuring: the three ground nodes in diagram b were combined into a single ground node, shown in the bottom of diagram c.

The fundamental feature emphasized by diagram c is that the two potential sources *ψ*_soil_(*t*) and *ψ*_leaf_(*t*) are clearly seen connected in parallel together with the capacitor branch. Here, we use branch in the electric sense, meaning the elements between two nodes, not the actual branch of a plant. For this reason, it is *not possible* to assume that a single effective potential difference ∆*ψ*(*t*) = *ψ*_soil_(*t*) *− ψ*_leaf_(*t*) is driving the flow; this could only be accomplished if the potential sources were in series (Alexander and Sadiku, 2012). The main conclusion is that *independent* potential sources are necessary whenever an internal water storage is present.

Another fundamental feature of the electric analogue is that the capacitor connects to the main line on one side, and it is grounded (i.e., connected to a zero potential) on the other side, as discussed in rule 4.

These two fundamental features of the electric analogue — independent potential sources and a grounded capacitor — have been overlooked by previous studies that took the electric approach (Landsberg et al., 1976; Jones, 1978; Milne et al., 1983; Dalton, 1995; Phillips et al., 1997; Nobel et al., 1999; Phillips et al., 2004; Zhuang et al., 2014). Some depict electric analogues with one source only, either a potential (voltage) source or a flow (current) source. In essence, a hidden assumption in these models is that the flow that leaves the plant towards an effective potential difference ∆*ψ* is the same flow that then enters the plant — a potential difference does not create flow, it only produces a potential step. A single potential difference ∆*ψ* in effect cancels any possibility of the internal water storage to contribute extra flow in case of increased evaporative demand, defeating the very purpose of the storage. Some of the models also violate the grounded capacitor feature, and they connect the capacitor twice to the main line. This means that the internal water storage unit in these models is implicitly being directly controlled by both the potential in the the plant’s xylem (*ψ*_*X*_, which was intended) and by the potential in the soil (*ψ*_soil_, not intended). A judicious construction of the electric analogue for plant hydraulics is of critical importance, because it is on the equations that arise from it that we derive conclusions on the dynamics of the system.

Finally, the model represented by the diagrams shown in Fig. 1 is a *minimal model*. This means that in modeling the hydraulics of a plant with internal water storage, either with the hydraulic or with the electric interpretation, one cannot dispense with any of the constituents shown in Fig. 1. The resistance *R*_1_ could not be dispensed with, because this would mean that the xylem potential *ψ*_*X*_ is equal to the soil potential *ψ*_soil_, decoupling the dynamics in the capacitor branch from the dynamics in the branch with *R*_2_. The same argument works for *R*_2_ being a necessary part of the model. Finally, the resistance *R*_*c*_ cannot be dispensed with, because this would mean that the potential on the capacitor *ψ*_*W*_ would respond instantaneously to any changes in the xylem potential *ψ*_*X*_. Fast changes in *ψ*_*W*_ would amount to arbitrarily high changes in the internal water content (recharge flow), which is not realistic (e.g., Jones, 1978; Sperry et al., 1998; Bonan et al., 2014; Xu et al., 2016). Hunt et al. (1991) hypothesized that the minimum number of constituents necessary to represent the water flow through a whole plant is one capacitor and one or two resistors. Our analysis shows that one capacitor and three resistors, arranged as shown in Fig. 1, would be the least one could do.

The system of equations that describes the dynamics of the minimal model is given by

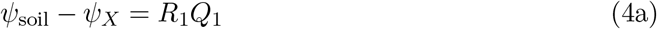

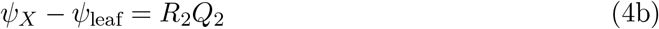

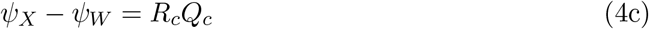

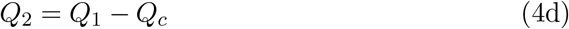

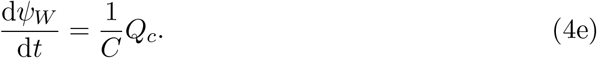

Equations (4a), (4b) and (4c) derive from Eq. (1) applied to resistances *R*_1_, *R*_2_ and *R*_*c*_, respectively (rule 1). Equation (4d) derives from flow conservation (rule 2) in the node labeled X, and Eq. (4e) derives from the time derivative of Eq. (3) in rule 4, where the recharge 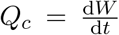. One can solve the five Eqs. (4) for the five unknowns *Q*_1_, *Q*_2_, *Q*_*c*_, *ψ*_*X*_, 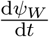, yielding one differential equation for *ψ*_*W*_

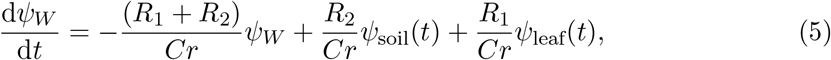

and four equations for the other unknowns,

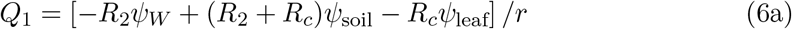

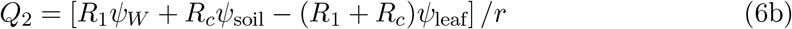

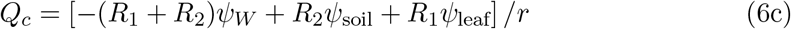

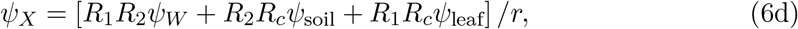

where *r* = *R*_1_*R*_2_ + *R*_1_*R*_*c*_ + *R*_2_*R*_*c*_. In order to know everything about the dynamics of our system, it suffices to solve Eq. (5) for *ψ*_*W*_ (*t*), and substitute the result in Eqs. (6).

A convenient way of solving the equations above for arbitrary forcing *ψ*_soil_ and *ψ*_leaf_ is provided by System Dynamics. The most important mathematical entity that fully captures the essence of our system, and that is unequivocally able to describe its dynamics, is the transfer matrix ***G***(*s*),

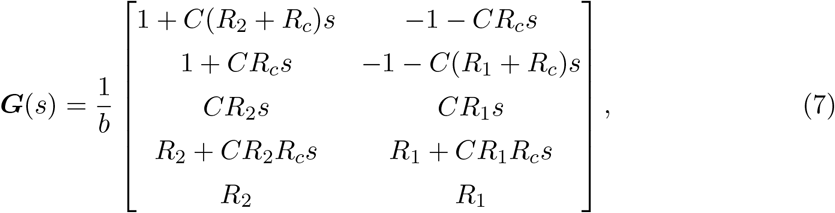

where *b* = *R*_1_ + *R*_2_ + *Crs*, and *s* in the complex frequency. As a rule, the mathematical derivations for the expressions used in this paper can be found in the Supporting Information (SI). For the derivation of ***G***(*s*) see SI.1. This matrix has five lines, each corresponding to five unknowns (*Q*_1_, *Q*_2_, *Q*_*c*_, *ψ*_*X*_, *ψ*_*W*_), and two columns, each corresponding to a different forcing (*ψ*_soil_, *ψ*_leaf_). If, for instance, we would like to know how transpiration *Q*_2_ (row 2) responds to changes in *ψ*_soil_ (column 1), we need to appropriately examine the matrix element ***G***_21_(*s*).

In the next section we will use the tools of system dynamics, in particular analyzing the transfer matrix ***G***(*s*), to gain insight into the role of the internal water storage in the dynamics of flows and water potentials throughout the plant.

## Results

### Transient response to step forcing

We investigate how a plant in steady state responds to sudden changes in the drivers of the dynamics, in this case the soil and leaf water potentials (*ψ*_soil_, *ψ*_leaf_). A step change to either of these potentials will bring the plant to a new steady state, and the transient response of the water flows (*Q*_1_, *Q*_2_, *Q*_*c*_) is uniquely determined by the plant traits (*R*_1_, *R*_2_, *R*_*c*_, *C*).

Figure 2 shows the dynamics of the water flows *Q*_1_ and *Q*_2_ for two cases, where either *ψ*_soil_ (panel a) or *ψ*_leaf_ (panel b) are discontinuously changed. Both cases start with the same steady state, where constant 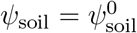 and 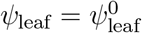 result in no recharge (*Q*_*c*_ = 0), and in constant and equal sap and transpiration flows *Q*_1_ = *Q*_2_ = ∆*ψ/*(*R*_1_ +*R*_2_), where ∆*ψ* = *ψ*_soil_ *− ψ*_leaf_. These solutions are obtained by solving Eq. (5) for steady state, and then using (6a), and (6b). At time *t* = 0, ∆*ψ* is instantaneously increased by *A* = 1 MPa, resulting in new steady-state flows that are higher by *d* = *A*/(*R*_1_ + *R*_2_) with respect to the previous values.

**Figure 2:**
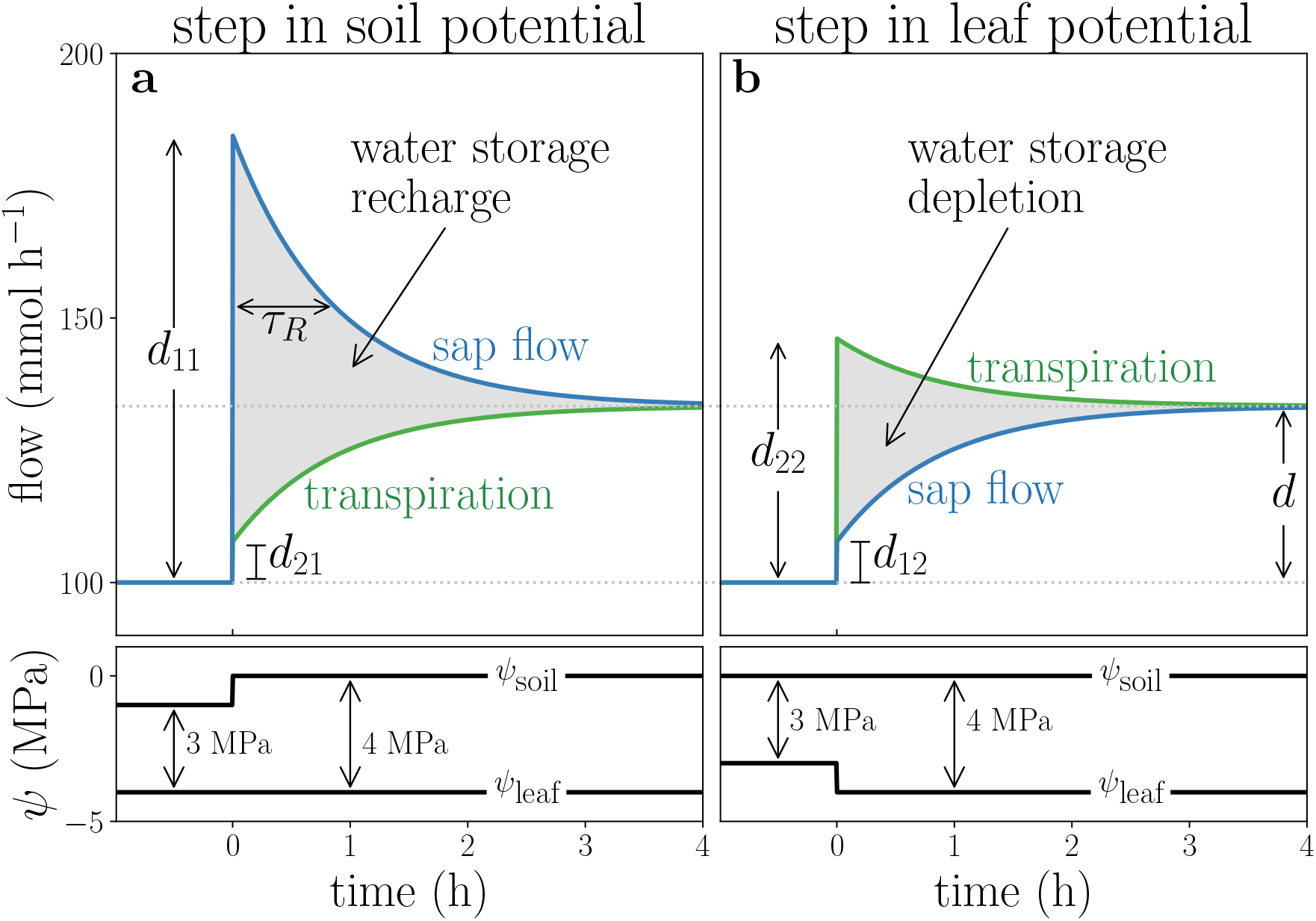
Step change in different environmental forcing results in opposite and non commensurate trends in flow dynamics. Panel a: sap flow exceeds transpiration flow (top), following a step change in soil water potential *ψ*_soil_ (bottom). Panel b: transpiration flow exceeds sap flow (top), following a step change in leaf water potential *ψ*_leaf_ (bottom). Parameter values: *R*_1_ = 0.01, *R*_2_ = 0.02, *R*_*c*_ = 0.002 (MPa h mmol^*−*1^), *C* = 100 (mmol MPa^*−*1^).

Although in both cases the step change in ∆*ψ* is exactly the same (from 3 to 4 MPa), the transient behavior of the flows is different, illustrating our previous assertion that ∆*ψ* can not be considered the driver of the dynamics, and that soil and water potentials must be treated separately. In the first case, depicted in panels a on the left, the increase in ∆*ψ* is due to an increase in *ψ*_soil_, and *ψ*_leaf_ is kept constant, while the opposite is true in panels b on the right, where *ψ*_leaf_ decreases and *ψ*_soil_ is kept fixed.

In case (a), sap flow will discontinuously increase by *d*_11_, always overshooting the steady state (*d*_11_ *> d*), while transpiration will increase by *d*_21_, which is always smaller than *d*. In case (b) the roles are reversed: the transpiration increases by *d*_22_, always overshooting the steady state (*d*_22_ *> d*), and sap flow will increase by *d*_12_. Values for *d*_*ij*_ are detailed in SI.2. The instantaneous increase in sap flow at the onset of transpiration (case b) supports Burgess and Dawson’s (2007) hypothesis that a cohesion-tension framework (like ours) would predict “small flows at the stem base commencing simultaneously with flows in the branches”. Indeed, as they suggest, this sap flow can be quite small and difficult to measure, since *d*_12_ = *AR_c_/r* can be much smaller than *d*.

The variation in water storage, which is the area between the two curves, is positive (storage recharge) for a positive step in soil potential (panel a), and it is negative (storage depletion) for a negative step in leaf potential (panel b). Although there seems to be a symmetry between the two cases because of the same change in ∆*ψ*, the volume of water storage depletion/recharge is not the same. In SI.2 we show that the ratio between the recharge volume of case a and the depletion volume of case b is *R*_2_/*R*_1_. For the parameter values in Fig. 2 this ratio is 2, meaning that the water storage in this case is twice as sensitive to a step change in soil potential then in leaf potential.

The characteristic time scale — relaxation time *τ*_*R*_ — under which the system responds to step-like forcing is given by

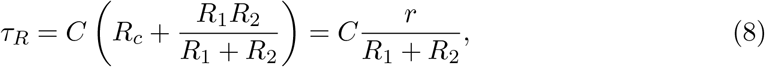

see SI.2. It bears emphasizing that this relaxation time scale *τ*_*R*_ is the only characteristic time scale of our system, and it applies to the dynamics of all quantities.

The expression above for *τ*_*R*_ differs from previous results in a few ways. Phillips et al. (1997) derived two time scales (for the ‘capacitive pathway’ and ‘total network’) for an electric diagram that does not conform to the fundamental features discussed before. In SI.3 we show how to reconcile their approach with ours. Hunt et al. (1991) and Wronski et al. (1985) provide an expression that is equivalent to ours, for the specific case that *R*_1_ = *R*_2_, i.e., the representative locus of the internal water storage is such that resistances to flow below and above this point are exactly the same.

To sum up: we have shown that the transient response of sap, transpiration and recharge flows discriminate between step changes in *ψ*_soil_ and *ψ*_leaf_, and that the effects of increasing soil potential are not the same as decreasing leaf potential by the same amount. Furthermore, we derived the expression for the relaxation time scale *τ*_*R*_, based on an accurate analogue diagram for the plant hydraulics.

### Frequency response to periodic forcing

The periodic change in soil or atmospheric conditions is an important and realistic situation that plants encounter, and we will now investigate how plant flows respond to it. When subjected to sinusoidal forcing of *ψ*_soil_ or of *ψ*_leaf_, our system settles in a periodic steady state with period equal to that of the forcing period. Since fluctuations in atmospheric conditions are usually much stronger than fluctuations in soil water, we will focus on the case of fixed *ψ*_soil_ and a varying leaf water potential, according to

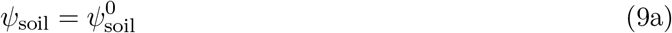

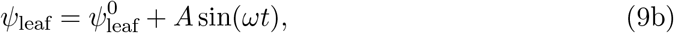

where *A* is the amplitude of the forcing, *ω* = 2*π/T* is the forcing frequency, and *T* is the forcing period. The analysis of the opposite case (fluctuating *ψ*_soil_ and constant *ψ*_leaf_) will be alluded to when necessary.

#### Flow amplitude response

How will the amplitude of the flows *Q*_1_, *Q*_2_, and *Q*_*c*_ respond to the driving force shown in Eq. (9)? Figure 3 (panels a–c) shows the response of these flows as a function of time, for three forcing periods of increasing length (8, 32, and 128 hours, the darker the shade, the longer the period). Increasing forcing frequency *ω* has the effect of increasing the amplitude of recharge, while sap flow amplitude decreases, and transpiration amplitudes stay approximately the same for all frequencies.

**Figure 3:**
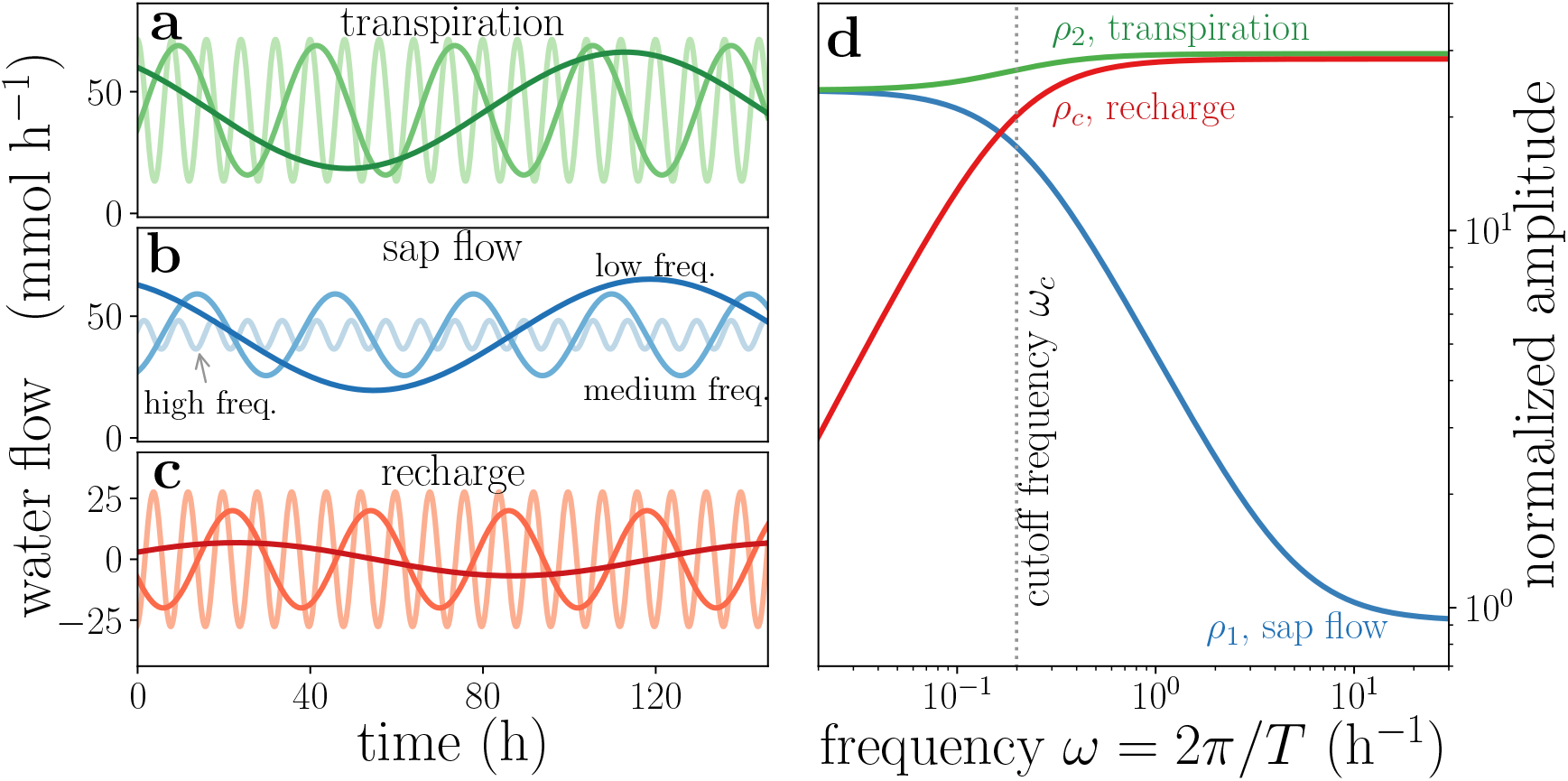
Plant behaves as a frequency filter, damping high frequencies from sap flow and low frequencies from recharge, while transpiration is mostly unaffected. Panels a–c: periodic dynamics of transpiration, sap flow, and recharge, for driving *ψ*_leaf_ of period 8, 32, and 128 hours (light, medium, and dark shades, respectively). Panel d: normalized amplitude of oscillation as function of angular frequency *ω*. Dotted line indicates cutoff frequency. Parameter values: *R*_1_ = 8.8 *×* 10^*−*3^, *R*_2_ = 3.4 *×* 10^*−*2^, *R*_*c*_ = 2.8 *×* 10^*−*4^ (MPa h mmol^*−*1^), *C* = 693 (mmol MPa^*−*1^).

The flows shown in Fig. 3a-c are described by

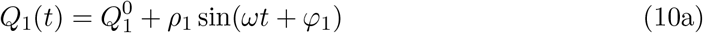

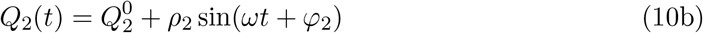

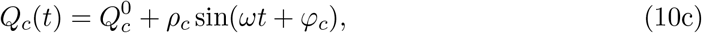

where *ρ* is the amplitude of oscillation, and *ϕ* is the phase. The oscillation in these flows occur around 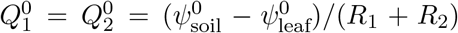 and 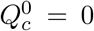, same as the steady-state values in the previous analysis. How does the amplitude of oscillation depend on the forcing frequency? Figure 3d shows the normalized amplitudes of oscillation (*ρ*_1_/*A*, *ρ*_2_/*A*, *ρ*_*c*_/*A*) as a function of the forcing frequency *ω*. The expressions for the amplitudes of oscillation and their detailed derivation are in SI.4. The decreasing curve for *ρ*_1_ is characteristic of a low-pass filter, where high frequencies are filtered out of sap flow *Q*_1_, as is also seen in panel b. Conversely, the rising curve for *ρ*_*c*_ is typical of a high-pass filter, where low frequencies are dampened from recharge *Q*_*c*_. A useful measure of the qualitative change in the frequency filtering is the cutoff frequency *ω*_*c*_, located at the “elbow” of the curves for *ρ*_*c*_ in panel d. The cutoff frequency for recharge *Q*_*c*_ is given by

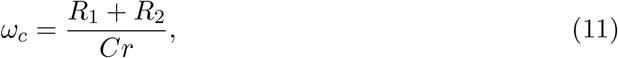

(see SI.4). For the parameter values of Fig. 3, *ω*_*c*_ ≃ 0.2 h^*−*1^, shown by the dashed vertical line in panel d. One can say, in an approximate manner, that frequencies above *ω*_*c*_ are mainly filtered out from *Q*_1_, while frequencies below *ω*_*c*_ are filtered out from *Q*_*c*_. Because the period associated with the cutoff frequency in this example is 2*π/ω*_*c*_ ≃ 32 hours, most of the daily fluctuations in atmospheric conditions are filtered from the sap flow response, and the faster the fluctuation, the greater the filtering. For all values of the parameters, the amplitude of the transpiration *ρ*_2_ will always be greater than that of sap flow *ρ*_1_ or the recharge *ρ*_*c*_ (see SI.4).

The filtering effect described here explains quantitatively why sap flow is always smoother than the transpiration signal (e.g., see Fig. 6). This filtering also means that a pulse in transpiration of a given strength and duration spreads as it moves down the plant. The transpiration signal, stripped of its highest frequencies, translates into shorter and wider sap flow pulses, because of the buffering granted by the internal storage.

The curve for *ρ*_2_ in panel d indicates that transpiration is able to readily respond to fluctuations of all frequencies, slow or fast. This means that the plant is able to maintain a steady transpiration flow, no matter the frequency of the forcing. For lower frequencies, transpiration flow *Q*_2_ is mostly supplied by sap flow *Q*_1_, and the internal water storage does not play an important role. The situation is reversed for higher frequencies, where transpiration is mostly supplied by the internal water storage, and not directly from the sap flow.

The results above regarding frequency filtering hold true not only for a sine-like forcing of *ψ*_leaf_, but for *any signal*, since it can always be decomposed into a sum of sines of various frequencies. The general result here is that, given the basic plant traits (*R*_1_, *R*_2_, *R*_*c*_, *C*), we can quantify the degree in which fast and slow environmental changes propagate and are dampened throughout the plant.

We assumed that only *ψ*_leaf_ varies, while *ψ*_soil_ was kept constant. Because of the symmetries of the model, assuming fixed *ψ*_leaf_ and sinusoidal changes in *ψ*_soil_ yields exactly the same results with indices 1 and 2 interchanged. For instance, the transpiration *Q*_2_ would now behave as a low-pass filter, but the behavior of *Q*_*c*_ would still be characteristic of a high-pass filter.

We can reinterpret the effects of step forcing seen before in light of the filtering properties of the system. A discontinuous jump in leaf water potential is composed of all frequencies (the Fourier transform of a step function is proportional to *ω*^−1^), but the higher frequencies will be filtered out of the sap flow. This is why transpiration readily responds to a step change in *ψ*_leaf_ in Fig. 2a (no significant filtering occurs), but sap flow, without the higher frequencies, evolves in a much smoother trajectory.

The daily contribution of internal water storage to transpiration can also be found by analyzing the flow amplitudes. This relative contribution is the ratio between the positive recharge over a day *T*_day_*ρ*_*c*_/*π*, and the mean daily transpiration 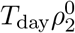 (see SI.4D), yielding

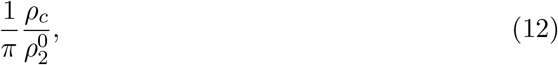

where *T*_day_ is the length of a day, and *ρ*_*c*_ [see Eq. (12c) in SI.4A] needs to be computed for *ω* = 2*π/T*_day_.

### Hydraulic safety margin

The buffering effect offered by the internal water storage can play an important role in preventing xylem water potential *ψ*_*X*_ from reaching very low values, which are associated with embolism and serious risk of hydraulic failure. Assuming again a periodic forcing on the leaf water potential only, given by Eqs. (9), the solution for *ψ*_*X*_ (*t*) [given by Eq.(6d)] will also respond periodically, oscillating sinusoidally around a mean value 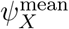 with an amplitude *A*_*X*_. Figure 4a shows three realizations of *ψ*_*X*_ (*t*), for three capacitance *C* values, and a forcing period of 24 hours.

**Figure 4:**
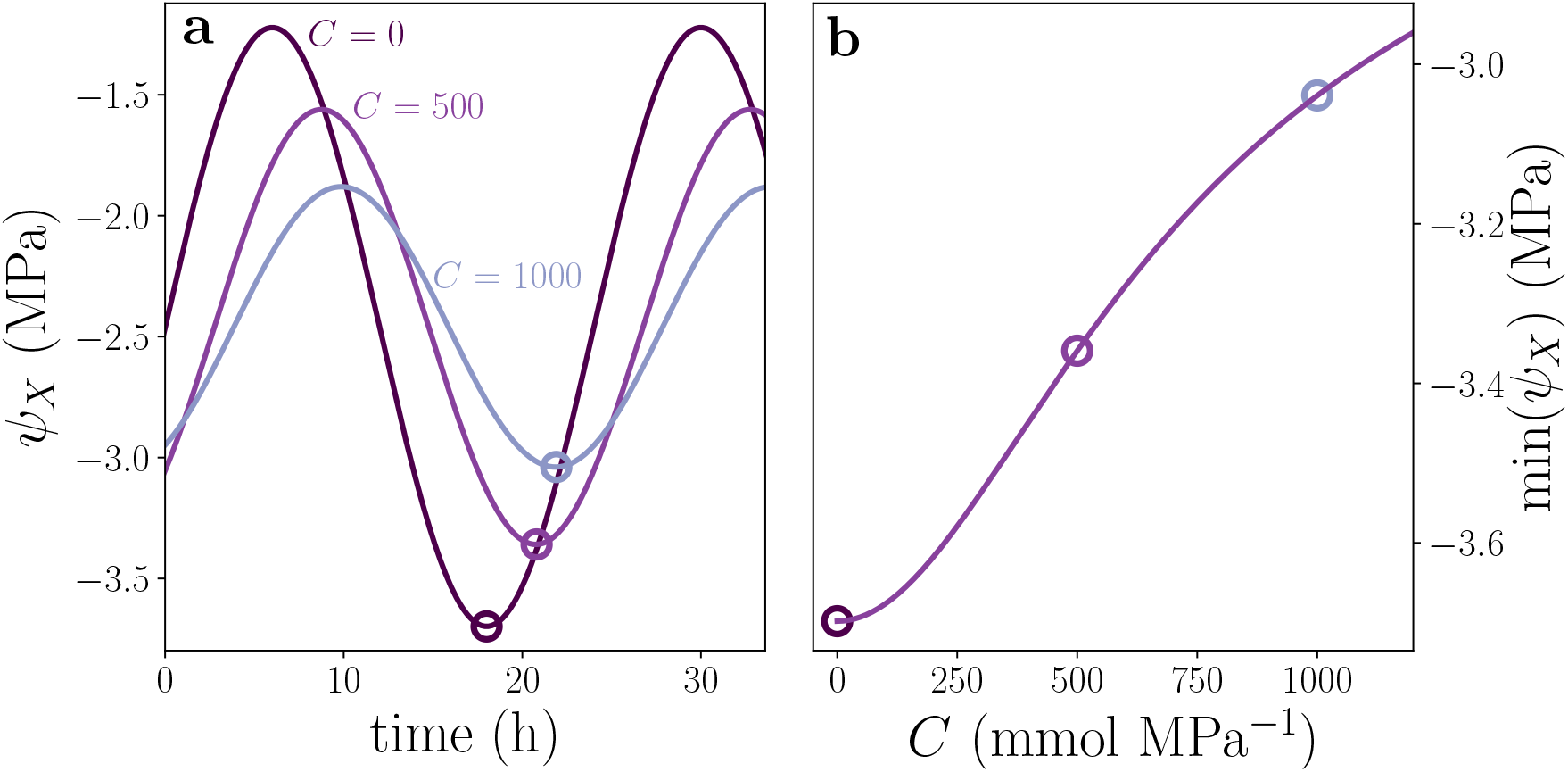
Minimal value of xylem water potential *ψ*_*X*_ increases with capacitance. Panel a: evolution in time of *ψ*_*X*_ for three capacitance values. Panel b: higher capacitance yields greater minimal *ψ*_*X*_ values. Parameters: same as in Fig. 3.

As the capacitance increases, the oscillation amplitude decreases. As a consequence of this, the minimal value 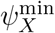 goes up for increasing *C*, see hollow circles in Fig. 4a. This means that the internal water storage confers the plant a hydraulic safety margin (Meinzer et al., 2009), helping to protect the plant from low xylem water potentials, thus decreasing the chance of embolism and an accompanying loss in hydraulic conductivity. Figure 4b shows the increase in 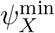 with higher capacitance values, where the expression for 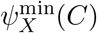 can be found in SI.5. This prediction is consistent with Meinzer et al. (2009, see Fig. 5a therein).

**Figure 5:**
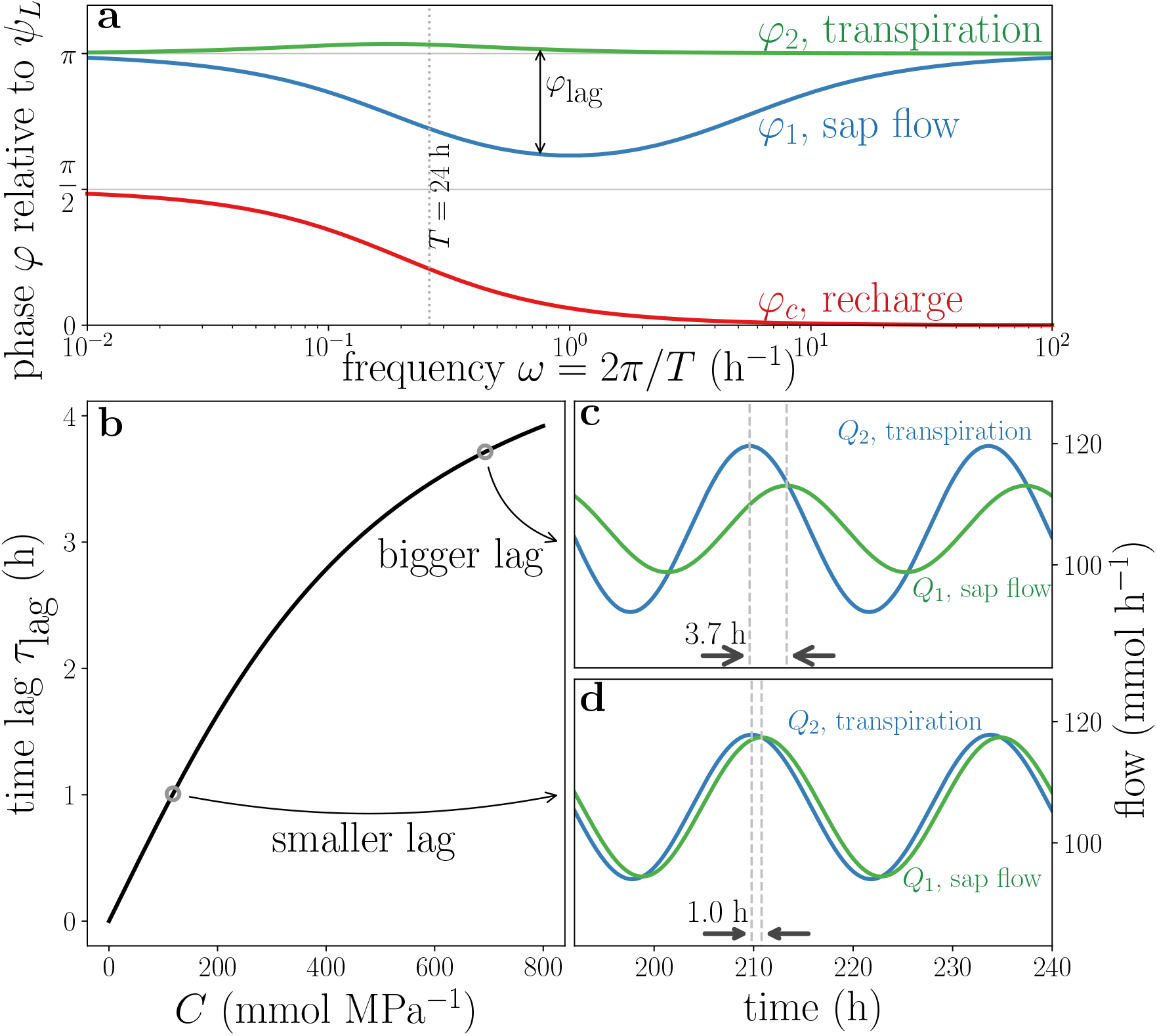
Time lag between *q*_2_ and *q*_1_ signal increases with capacitance. Panel a: Phase *ϕ* of the flows as function of angular frequency *ω*. Zero (*π*) phase denotes that flow is in phase (antiphase) with driving *ψ*_leaf_ signal. Panel b: Time lag *τ*_lag_ as a function of the capacitance *C*. Panels c and d: two realizations of the dynamics of sap flow and transpiration, for relatively big and small values of capacitance. Vertical dashed lines and thick arrows help emphasize the time lag between peak transpiration and peak sap flow in daily dynamics. Parameters: same as in Fig. 3.

### Phase and time lags

Not only the amplitude of oscillation of the flows (*Q*_1_, *Q*_2_, *Q*_*c*_) are influenced by varying forcing frequency *ω*, but also their phases *ϕ*. These phases [see Eqs. (10)] convey information on how much the flows are delayed or ahead of the forcing signal *ψ*_leaf_.

Figure 5a shows *ϕ*_1_, *ϕ*_2_, and *ϕ*_*c*_ as functions of the forcing frequency *ω*. We see that *ϕ*_2_ is in the vicinity of *π*, which means that when *ψ*_leaf_ is lowest (highest evaporative demand) transpiration *Q*_2_ will be at its highest approximately at the same time. Because *ϕ*_2_ is always slightly higher than *π*, the transpiration peak will be a bit before the minimum of *ψ*_leaf_.

Conversely, *ϕ*_1_ is always smaller than *π*, meaning that sap flow *Q*_1_ will peak after the minimum of *ψ*_leaf_. The phase lag *ϕ*_lag_ = *ϕ*_2_ *− ϕ*_1_ between these two flows means that sap flow *Q*_1_ will always lag behind transpiration *Q*_2_, delayed by a time lag *τ*_lag_, given by

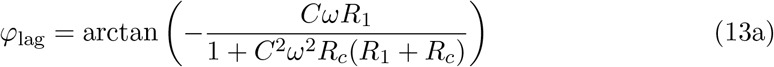

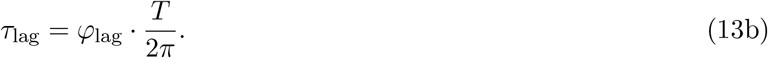

Details on the derivation of the phases and on the time lag are found in SI.6.

It is only because the plant has an internal water storage that transpiration can become decoupled from sap flow: if there were no water storage (*C* = 0), the time lag would be zero. This possibility of being momentarily in “hydraulic overdraft” during periods where transpiration is greater than sap flow could have a decisive role in the survival of plants under drought stress.

It is interesting to note that the expression for *ϕ*_lag_ does not depend on *R*_2_. An intuitive explanation for this is that *Q*_1_ lags behind *Q*_2_ because of the system constituents below the upper branch in Fig. 1a, namely *R*_1_, *R*_*c*_, and *C*. The signal from *ψ*_leaf_ reaches these constituents exclusively *after* it has passed the upper branch, through *R*_2_. Therefore, any further time lag in the *Q*_1_ signal with respect to *Q*_2_ can only be affected by *R*_1_, *R*_*c*_, *C*, but not by *R*_2_. This property will be useful in interpreting the results from the next section, where we evaluate our model.

Daily variations (*T* = 24 h) in *ψ*_leaf_ are of special importance: what can we say about the dependence of the time lag *τ*_lag_ on the plant traits? Figure 5b shows that *τ*_lag_ increases with capacitance *C*, for a forcing period of 24 h, according to Eqs. (13). Greater values of *C* translate into greater time lags, which can be seen in panels c and d, showing the response of the flows for high and low values of *C*, respectively (Hunt and Nobel, 1987).

### Parameterization and model evaluation

In this section we will see how much of the daily dynamics in plant hydraulics our model can capture. The only mechanism incorporated into the minimal model (besides the trivial Darcy-like saturated flow) is the internal water storage, and we left aside major mechanisms, for instance, stomatal control on transpiration. To the extent that this model captures certain behaviors—and fails to capture many others—, we gain insight on the role of the internal water storage.

We parametrized and evaluated our model against measurements in Yatir forest, a semi-arid pine plantation (280 mm mean annual precipitation), on the northern border of the Negev desert in Israel (Grünzweig et al., 2003; Rotenberg and Yakir, 2010). The available data was: eddy-covariance-based evapotranspiration flux (ET), sap flow (SF), stem diameter, soil-water content, air temperature, and relative humidity (Klein et al., 2014; Tatarinov et al., 2016). Because the period in question is towards the end of a six-month long dry season (September), ET is almost exclusively explained by transpiration (Rohatyn et al., 2018; Qubaja et al., 2020).

Figure 6 shows the measured ET, SF and stem diameter (a measure of change in stem water storage), all rescaled in order to emphasize the timing of their peaks. We optimized our model parameters for the ET data using the “Fitness Scaled Chaotic Artificial Bee Colony” algorithm, implemented by Python’s SPOTPY package (Houska et al., 2015). The optimal parameter values obtained are *R*_1_ = 8.8 *×* 10^*−*3^, *R*_2_ = 3.4 *×* 10^*−*2^, *R*_*c*_ = 2.8 *×* 10^*−*4^ (MPa h mmol^*−*1^), *C* = 693 (mmol MPa^*−*1^), and were used in Figs 3, 4, and 5. ET (green line) peaks in mid-morning and in late afternoon, showing a typical midday depression in transpiration, while stem diameter and SF peak, respectively, before and after the major peak in ET. The arrows on the top show the time lag between maximum ET and maximum stem diameter, while the arrows on the bottom show the time lag between ET and SF.

**Figure 6:**
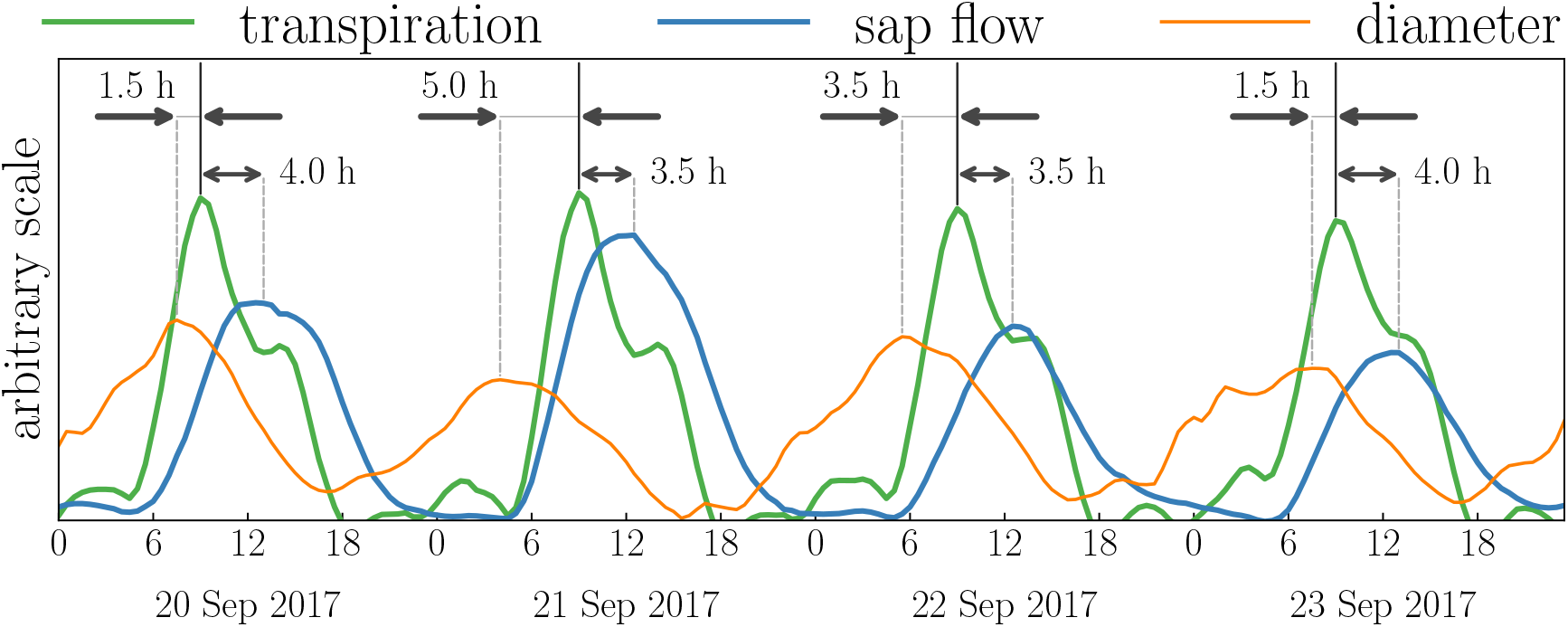
Measured evapotranspiration (ET), sap flow (SF), and tree diameter in a semi-arid pine forest, over four days during the dry season. The arrows on the top indicate time lag between ET and stem diameter, while arrows on the bottom show time lag between ET and SF. Modeled time lag between ET and SF is shown in Fig. 5c.

Thanks to the estimation of optimal values for our model’s parameters, we can now make use of many of the predictions yielded by our model and see how they perform against measured data. These parameters condense information on whole-plant traits related to water storage (*C*) and hydraulic resistance (*R*_1_, *R*_2_, *R*_*c*_).

The cutoff frequency calculated for the optimized parameters is *ω*_*c*_ = 0.2, meaning that variations in atmospheric conditions faster than 32 hours will be mostly damped from SF. The diel fluctuations are clearly found in SF, but there is no trace of ET midday depression on it, since the midday depression has a much shorter time scale (about 3 hours), and is thus filtered out.

The time lag between ET and SF averages 3.75 hours over the four-day period shown in the figure, while the predicted time lag, according to Eq. (13b), is *τ*_lag_ = 3.71 hours. Stomatal control, if introduced to our model, would effectively be expressed as a varying resistance *R*_2_, but that is precisely the one factor that does not contribute to *τ*_lag_, as discussed before. This showcases the strength of our modeling approach: some patterns in the plant water dynamics are strong enough that even a minimal model like ours is able to capture them.

Another time lag can be measured, that between the stem diameter and ET, shown by the arrows on the bottom of Fig. 6b, and averaging 2.9 hours over the four days. Assuming that the stem diameter has a linear relation with the amount of water in the plant (*W*), then according to Eq. (3) we have that stem diameter has a linear relation with *ψ*_*W*_. The expression for the time lag between *ψ*_*W*_ and *Q*_2_ is shown in SI.6 (it also does not depend on *R*_2_), and for the optimized parameters it gives 3.9 hours. Not only does the model yield a reasonable estimate for this time lag, the discrepancy is consistent with our expectation. The stem diameter is measured lower in the tree than the representative height where the internal water storage would be located, therefore the signal for stem diameter would be delayed with respect to the water potential of this internal storage.

Using Eq. (12) with the optimized parameters, we find that 31% of the daily transpiration was due to the internal water storage. Water balance of measured transpiration and sap flow for a similar period yields a figure of 35%.

## Discussion

In this paper, we used a minimal model for plant hydraulics to investigate the interplay between the environment and the internal water storage. We derived predictions regarding the time scales and magnitudes of important flows and water potentials in the plant. When evaluated against measurements, the model yields values for a low number of parameters that represent major whole-plant traits. These results, we believe, are helpful to recognize patterns and trends in the behavior of plants under drought stress. We characterized in detail two survival-enhancing effects of the internal storage: the hydraulic safety margin protecting xylem from embolism, and the possibility of a momentary “hydraulic overdraft” when transpiration is higher that root uptake flow. Because these survival-enhancing behaviors granted by the internal storage occur on sub-daily time scales, we believe that transient descriptions are warranted in understanding and predicting plant fluxes in drought-stressed ecosystems.

The minimal model is general, and the basic behavior it shows is valid for plants of different species and sizes, given appropriate parametrization. Indeed, this model can also be understood to represent the collective behavior of a number of individuals, not necessarily identical. In this case, the parameters would represent whole-plot or ecosystem traits, averaged over individuals that share the same soil and atmospheric conditions. When high-frequency flux data are available, this model can help calibrate plant parameters averaged over the large areas represented by individual pixels in regional and general circulation models.

On the other side of the size spectrum, the model presented here can be also understood to describe the dynamics of specific plant parts, such as the stem or leaves. One can stack a number of the minimal structures into layers, and obtain more refined parsing of the plant hydraulics (e.g., Cowan, 1972; Nobel and Jordan, 1983; Hunt et al., 1991; Xu et al., 2016). For instance, the results shown here can provide insight into the role of the hydraulic capacitance of leaves in rapidly supplying water for transpiration, while buffering oscillations in leaf water potential.

We based the model on realistic descriptions of water flow and capacitive storage, although not in their full complexity. Given our goal of understanding the most fundamental processes in plant hydraulics, which steps can be taken in order to expand the model’s predictive power? First and foremost, a simple mechanism for stomatal control would help elucidate which aspects of daily transpiration are due to the internal water storage, the stomatal regulation, or the interaction between these processes. Indeed, including a stomatal control mechanism would make the leaf water potential *ψ*_leaf_ as an internal variable of the model, and vapor pressure deficit would now be the external driver. Leaf potential *ψ*_leaf_ would now be granted a hydraulic safety margin because of the internal storage, in the same way as described for xylem water potential *ψ*_*X*_. This safety margin in *ψ*_leaf_ would have far-reaching consequences in stomatal regulation: the internal storage would not only provide readily-accessible water volume for transpiration, it would enable stomata to stay open for longer when evaporative demand is high.

In the model development, we also left aside nonlinearities in the flow due to embolism, and in the capacitance due to nonlinear pressure-volume relations in the tissues that hold the water storage. Thanks to that, we were able to fully solve a linear system with the tools of system dynamics. The inclusion of these nonlinearities would have quantitative effects on our predictions, but qualitatively, the phenomena described would be unchanged. For instance, a non-constant capacitance would change the value of the time lag between transpiration and sap flow, but the fundamental understanding of why sap flow lags behind transpiration would still hold. The patterns in plant hydraulics described here can serve as a roadmap, indicating to more detailed (and realistic) models where to focus their attention. On the other hand, the detailed models are indispensable in delineating the validity limits of conclusions derived from simpler models. This dialogue between modeling approaches is essential for a full account of plant and ecosystem functioning in all its richness.

## Acknowledgements

This research was supported by the Ring Family Foundation. We would like to thank Dan Yakir for his support and guidance. We would also like to thank Gabriele Manoli, Eli Rivkin, Michael Margaliot, and Vitaly Shaferman for useful discussion.

## Author contributions

AK and YM built the model, analyzed it, and wrote the first versions of the manuscript. YP and JMG designed and performed the experiments, and contributed to the final version of the manuscript.

## Supporting Information

### 1 System dynamics

Equations (5) and (6) in the main text form a linear and time-invariant system: all the expressions depend linearly on the dynamical variable *ψ*_*W*_ and on the inputs *ψ*_soil_, *ψ*_leaf_, and the coefficients do not depend on time. We can rewrite these equations in vector form (Ogata, 2004):

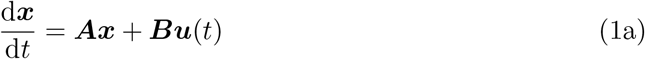

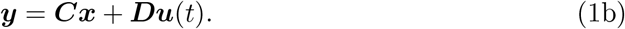

The state vector ***x*** = [*ψ*_*W*_] is a 1 *×* 1 vector in our case, and in general it is of size *n ×* 1, where *n* is the number of state variables, or the number of first-order differential equations to be solved. The 2 *×* 1 input vector ***u*** = [*ψ*_soil_, *ψ*_leaf_]^*T*^ denotes all the external influences on the system (it is of size *r ×* 1 for *r* inputs), and the 5 *×* 1 output vector ***y*** = [*Q*_1_, *Q*_2_, *Q*_*C*_, *ψ*_*X*_, *ψ*_*W*_]^*T*^ includes all information about which we would like to know the dynamics (in general of size *m×* 1 for *m* outputs). The output vector can contain any information we wish to know about the system, so in addition to the four unknowns shown in Eq. (6), we added *ψ*_*W*_ to the list. The matrices ***A**, **B**, **C**, **D*** are respectively called state matrix (size *n × n*), input matrix (*n × r*), output matrix (*m × n*) and direct transmission matrix (*m × r*), and are given by

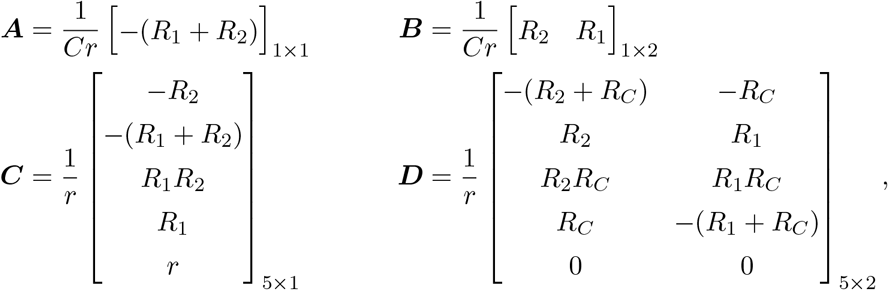

where the subscripts indicate the dimension of the matrices, in rows *×* columns.

The general problem of solving the linear and time-invariant system of Eq. (1) for arbitrary external input ***u***(*t*) can be accomplished by using the Laplace transform, that converts differential equations with respect to time *t* into algebraic equations with respect to the complex frequency *s*. The quantities ***y*** we wish to find are thus given by

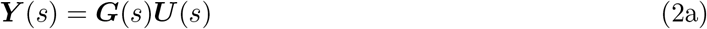

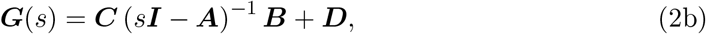

where ***Y*** (*s*) and ***U*** (*s*) are the Laplace transform of ***y***(*t*) and ***u***(*t*), and ***I*** is the identity matrix. Substituting the expressions for ***A**, **B**, **C**, **D*** into Eq. (2b) yields the transfer matrix ***G***(*s*):

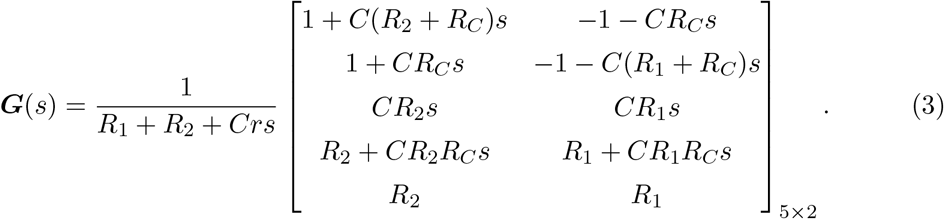

### 2 Step forcing

#### Relaxation time scale *τ*_*R*_

The characteristic time scale of a given variable (output) *i* and forcing (input) *j* is given by the inverse of the value of *s*, for which the denominator of the transfer matrix element ***G***_*ij*_(*s*) equals zero. In the language of system dynamics, the time scales *τ*_*R*_ are the inverse of the poles of the transfer function, which are the same as the eigenvalues of matrix ***A***. All matrix elements of ***G***(*s*) have the same denominator, namely *R*_1_ + *R*_2_ + *Crs* [see Eq. (3)], a polynomial of degree 1. To find *τ*_*R*_ we need to solve *R*_1_ + *R*_2_ + *Cr*/*τ*_*R*_ = 0, which gives *τ*_*R*_ = *Cr/*(*R*_1_ + *R*_2_). There are no different time scales for transpiration, recharge, sap flow, etc: they all have the exact same *τ*_*R*_.

#### Size of discontinuous jumps

For *t ≥* 0, the flows *Q*_1_ and *Q*_2_ evolve according to

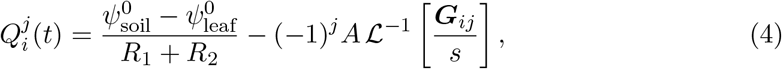

where the first term in the right-hand side is the steady state before the step change, the index *j* denotes the input that is being changed (*j* = 1 means *ψ*_soil_, *j* = 2 means *ψ*_leaf_), *−*(*−*1)^*j*^*A* accounts for positive/negative step changes, and 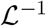 is the inverse Laplace transform. Rewriting ***G***_*ij*_ (for *i, j* = {1, 2}) in the Bode form

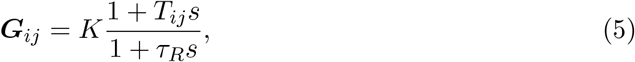

gives the solution

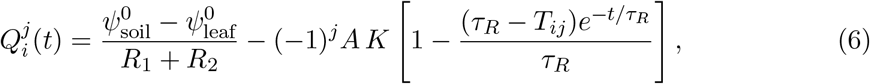

where *K* = (*R*_1_ + *R*_2_)^*−*1^, *T*_11_ = *C*(*R*_2_ + *R*_*c*_), *T*_12_ = *T*_21_ = *CR*_*C*_, and *T*_22_ = *C*(*R*_1_ + *R*_*c*_).

Therefore, for *t* = 0, the flows increase by

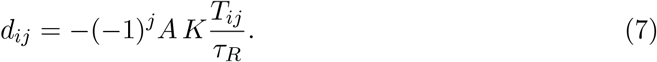

Substituting the relevant *T*_*ij*_, we find that the discontinuous jumps in *Q*_1_ and *Q*_2_ read

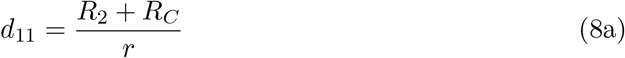

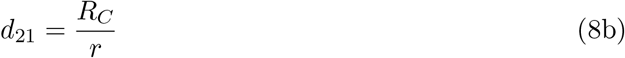

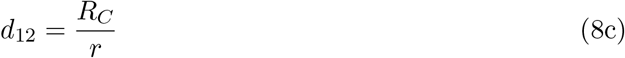

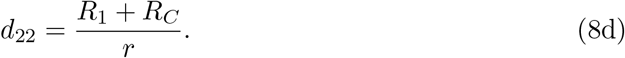

We know that *Q*_1_ overshoots for changes in *ψ*_soil_ and *Q*_2_ overshoots for changes in *ψ*_leaf_ because both *T*_11_/*τ*_*R*_ > 1, *T*_22_/*τ*_*R*_ > 1. Conversely, *Q*_2_ stays below the steady state for changes in *ψ*_soil_, and *Q*_1_ stays below the steady state for changes in *ψ*_leaf_ because *T*_12_/*τ*_*R*_ = *T*_21_/*τ*_*R*_ < 1.

#### Recharge/depletion ratio

The effect of a step change in *ψ*_soil_ or *ψ*_leaf_ on the water storage recharge flow 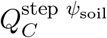 and 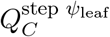 is calculated with the transfer matrix elements ***G***_31_ = *R*_2_*g* and ***G***_32_ = *R*_1_*g*, where *g* = *Cs/*(*R*_1_ + *R*_2_ + *Crs*). Because the Laplace transform of a step (Heaviside) function *H*(*t*) is simply 1/*s*, this ratio reads

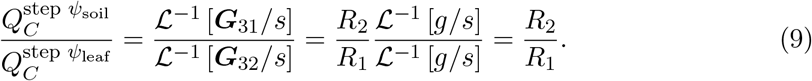

### 3 Thévenin equivalent

As shown in Fig. 1a, the potential sources *ψ*_soil_ and *ψ*_leaf_ are in parallel, and therefore we cannot combine them as ∆*ψ* = *ψ*_soil_ *− ψ*_leaf_, and assume that ∆*ψ* is driving the flow. However, there is a way to combine these two sources, by applying Thévenin’s theorem (Alexander and Sadiku, 2012) on the part of the circuit enclosed by a dotted rectangle. This conversion treats the left branch with the capacitor as the load of the circuit, thus giving it a special role in the dynamics.

**Figure 1:**
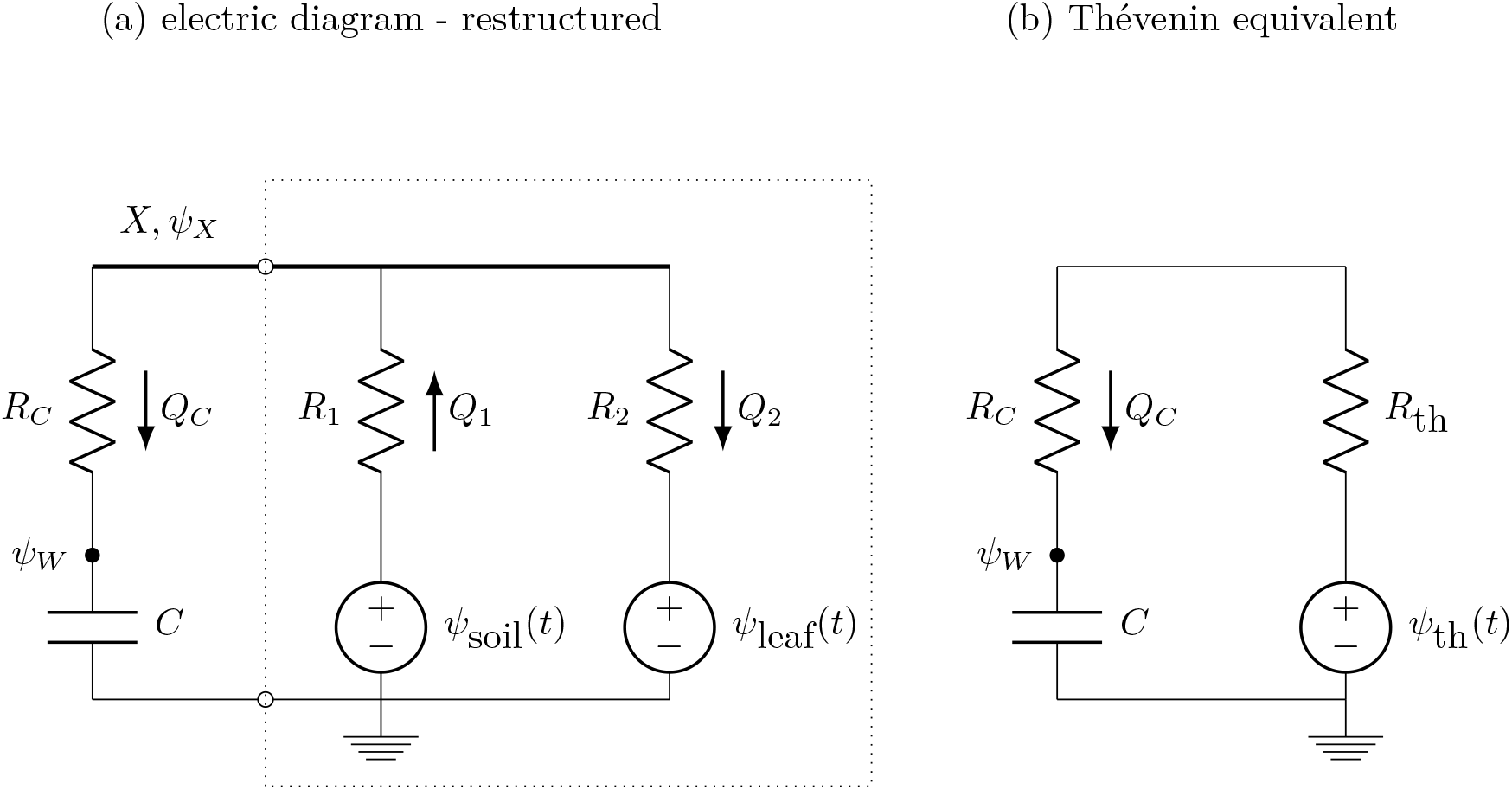
Clumping together the two water potential sources *ψ*_soil_ and *ψ*_leaf_ is possible, but it is not useful nor enlightening. Panel a shows the original electric diagram, and panel b shows its Thévenin equivalent. The components enclosed by the dotted rectangle on panel a were converted to equivalent Thévenin resistance and potential source. In the Thévenin equivalent diagram we can no longer talk about sap flow *Q*_1_ nor transpiration *Q*_2_.

All the resistances and potential sources in the dotted rectangle can be substituted by the Thévenin equivalent resistance and potential, given by

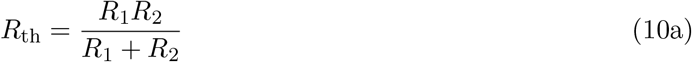

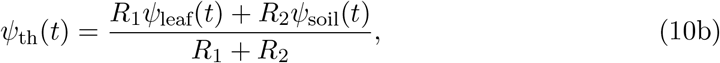

and we can now draw the equivalent circuit shown in Fig. 1b.

In the light of this conversion to the Thévenin equivalent circuit, the differential equation (5) in the main text for the dynamics of water storage potential simplifies to

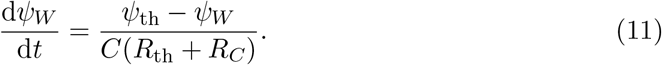

Equation (10b) shows that the soil and leaf water potentials can be combined attributing correct weights to them, and we can say that it is simply *ψ*_th_ that drives the dynamics. However, this statement would be true only considering the dynamics of the internal water storage. The conversion to the Thévenin equivalent eliminated from the dynamics important flows that we care about, namely the sap flow *Q*_1_ and transpiration *Q*_2_. These flows are nowhere to be found in Fig. 1b, limiting the use and insights to be gained by this simplified approach.

The results provided by Phillips et al. (1997) are based on such a simplified diagram, provided that, with the help of Norton’s theorem, their diagram showing a current source in parallel with a resistor is converted into the diagram in Fig. 1b (a voltage source in series with a resistor).

The significance of the discussion above is that modeling tree hydraulics with a system even simpler that our minimal model can be done, but necessarily such a model would yield partial information, and with parameters that are a nontrivial combination of parameters representing plant traits.

### 4 Periodic forcing: flow amplitudes

#### 4.1 Flow amplitudes

Expressions for the flow amplitudes under periodic forcing [Eqs. (9) in the main text] can be easily achieved, they are the absolute values of the relevant matrix elements ***G***(*iω*) (Ogata, 2004). For instance, *ρ*_*C*_/*A* = abs [***G***_32_(*iω*)], since *ρ*_*C*_ is the third element in the output ***y***, and *ψ*_leaf_ (the input that varies sinusoidally) is the second element in the input vector ***u***. The flow amplitudes, normalized by the amplitude of *ψ*_leaf_(*t*), read

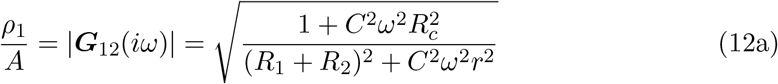

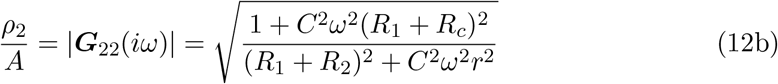

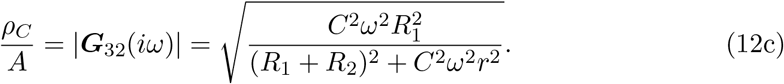

#### 4.2 Cutoff frequency

The cutoff frequency for *ρ*_*C*_ is the frequency for which *ρ*_*C*_ decreases by a factor of 1/*^√^*2 of its maximal value. This maximal value is

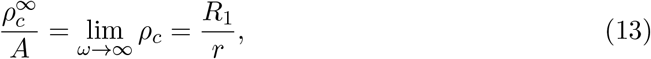

therefore solving 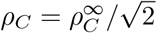 for *ω*_*c*_ yields

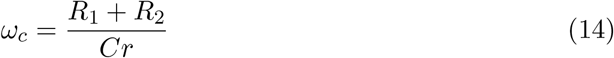

It is interesting to note that, for our simple system, the cutoff frequency *ω*_*c*_ is the inverse of the relaxation time *τ*_*R*_.

#### 4.3 Amplitude inequalities

The amplitude of transpiration *Q*_2_ is always greater than that of sap flow *Q*_1_ or recharge *Q*_*c*_:

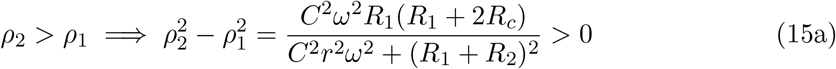

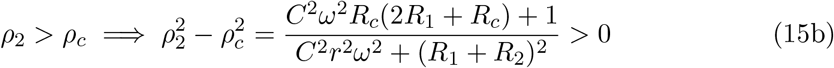

This can be seen visually in Fig. 3 in the main text: the green curve for *ρ*_2_ is always greater than the blue curve (*ρ*_1_) and than the orange curve (*ρ*_*C*_).

#### 4.4 Water storage contribution to transpiration

Recharge *Q*_*c*_ averages zero over a day (*Q*^0^), but the amount of water storage that contributes to daily transpiration is what leaves the storage over half a day:

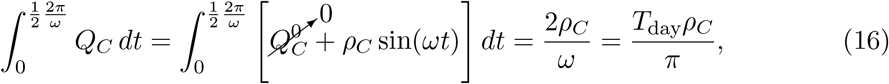

where the time translation corresponding to the phase *ϕ*_*C*_ was omitted for the sake of simplicity.

Daily transpiration is given by

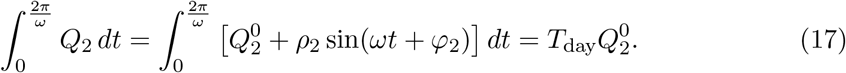

Therefore, the ratio *f* between daily water storage discharge and total daily transpiration is

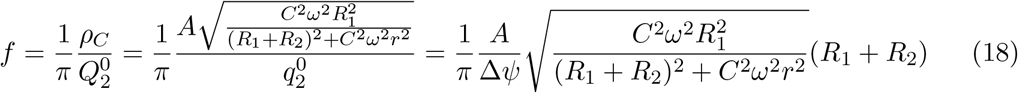

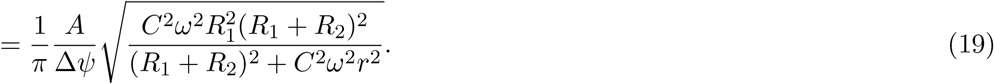

If we assume that *ψ*_leaf_ has its daily maximum equal to 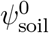 (leaf water potential equalizes with soil water potential), then we have that *A* = ∆*ψ*, and the ratio further simplifies to

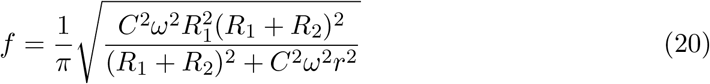

Figure 2 shows this fraction *f* as a function of the capacitance *C*, for resistance values obtained in the parametrization (values in Fig. 6). For the capacitance value in the optimized parameters, we have that the internal water storage represents 31% of total daily transpiration.

**Figure 2:**
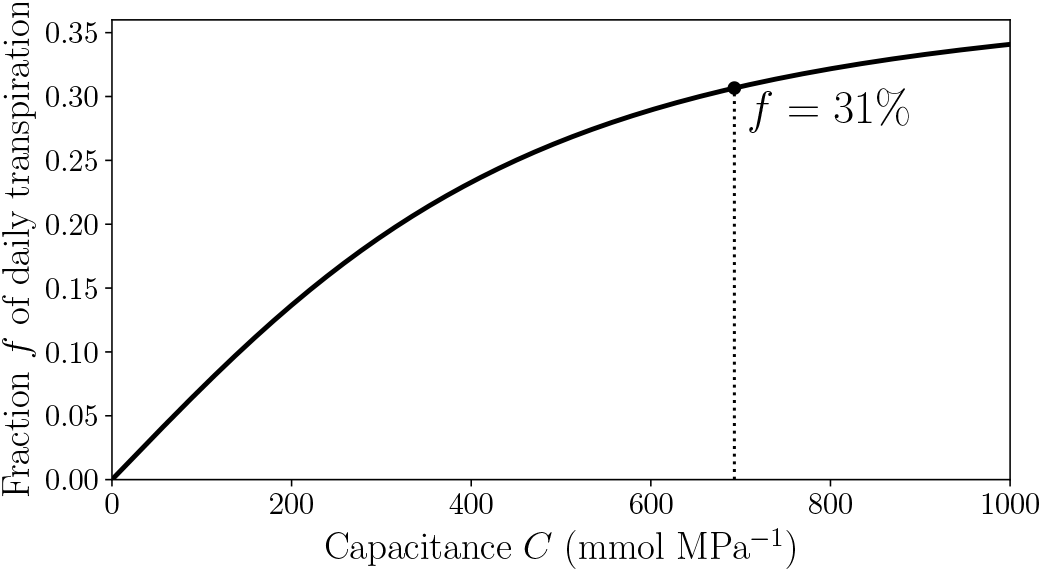
Fraction of daily transpiration that comes from internal storage increases with plant capacitance. The dependence of fraction *f* on capacitance *C* is shown in Eq. (20). For optimized parameters (see their values in Fig. 3 in the main text) the daily fraction is 31%.

### 5 Hydraulic safety margin

The minimal value assumed by *ψ*_*X*_ is given by

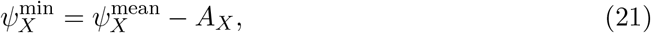

where the mean value 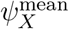 around which *ψ*_*X*_ oscillates can be found by solving Eq. (5) and (6d) in the main text, assuming steady state. The amplitude of oscillation *A*_*X*_ is given by *A ⋅* abs [***G***_42_(*iω*)], and A is the amplitude of oscillation in *ψ*_leaf_. Equation 21 can be rewritten as

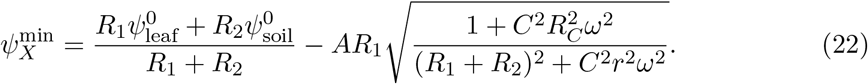

### 6 Phases *ϕ* and phase lag

The phases for *Q*_1_, *Q*_2_, and *Q*_*c*_ are given by

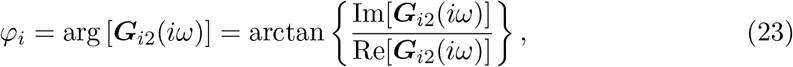

where *i* = 1, 2, 3 respectively. We have then

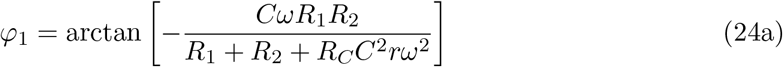

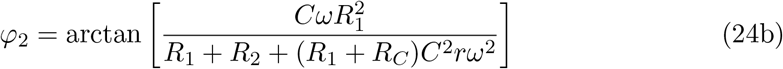

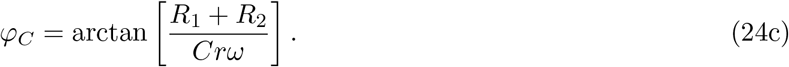

In order to calculate the phase lag *ϕ*_lag_ = *ϕ*_2_ *−ϕ*_1_, we can use the trigonometric identity

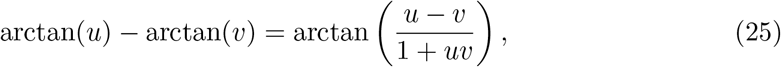

to yield

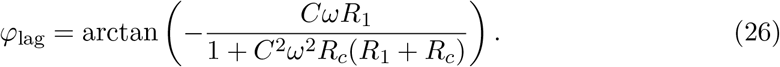

The phase lag between *Q*_2_ and *ψ*_*W*_ is similarly achieved:

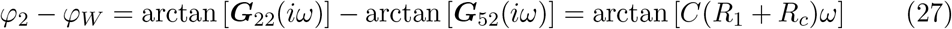

